# Efficient Multivariate Analysis Algorithms for Longitudinal Genome-wide Association Studies

**DOI:** 10.1101/394197

**Authors:** Chao Ning, Dan Wang, Lei Zhou, Julong Wei, Yuanxin Liu, Huimin Kang, Shengli Zhang, Xiang Zhou, Shizhong Xu, Jian-Feng Liu

## Abstract

**Motivation:** Current dynamic phenotyping system introduces time as an extra dimension to genome-wide association studies (GWAS), which helps to explore the mechanism of dynamical genetic control for complex longitudinal traits. However, existing methods for longitudinal GWAS either ignore the covariance among observations of different time points or encounter computational efficiency issues.

**Results:** We herein developed efficient genome-wide multivariate association algorithms (GMA) for longitudinal data. In contrast to existing univariate linear mixed model analyses, the proposed new method has improved statistic power for association detection and computational speed. In addition, the new method can analyze unbalanced longitudinal data with thousands of individuals and more than ten thousand records within a few hours. The corresponding time for balanced longitudinal data is just a few minutes.

**Availability and Implementation:** We wrote a software package to implement the efficient algorithm named GMA (https://github.com/chaoning/GMA), which is available freely for interested users in relevant fields.

## Introduction

Genome-wide association studies (GWAS) have been used to detect many genetic variants associated with various quantitative traits and complex diseases. Linear mixed models (LMM) adopted to GWAS (Kang, *et al.*, 2008; Lippert, *et al.*, 2011; *Yu, et al., 2006*; Zhou and Stephens, 2012) are able to capture genetic correlation among individuals, correct confounding environmental factors and control population stratification. However, most LMM based GWAS analytical tools, such as EMMA/EMMAX (Kang, *et al.*, 2010; Kang, et al., 2008), FaST-LMM (Lippert, et al., 2011), GEMMA (Zhou and Stephens, 2012) and GCTA (Yang, *et al.*, 2011), focus on traits that are measured only once. There are few methods available for GWAS dealing with longitudinal traits that are repeatedly measured during the life span of individual development.

Longitudinal traits, also known as dynamic traits or functional traits, are dynamically changing over a period of time controlled by both genetic effects and environmental factors. Multiple measurements at various time points during a life cycle are usually collected as longitudinal traits. Recently, advanced dynamic phenotyping system in animal and plant genetic experiments (Fahlgren, *et al.*, 2015; Porto, *et al.*, 2015) makes it feasible to acquire high throughput time varied datasets. Such repeated measurements under varying environmental conditions can improve statistical power of quantitative trait nucleotide (QTN) detection and help to further explore the mechanism of dynamical genetic control for complex longitudinal traits (Li and Sillanpaa, 2015; Wu and Lin, 2006). Analyzing such types of datasets also promotes early prediction of longitudinal traits and diseases (Kellogg, *et al.*, 2014; *McSweeney, et al., 2014*).

However, currently employed analytical methods, such as varying-coefficient regression (Gong and Zou, 2012) and estimation equation (Xiong, *et al.*, 2011), are computationally intensive compared to the univariate counterpart. An alternative way to improve computational efficiency is to analyze each single time point separately and then integrate test statistics across time points to determine the overall significance (Kwak, *et al.*, 2014). However, the single time point analysis is inefficient in QTN detection because it ignores the covariance among observations of different time points.

Random regression models (RRM) are multivariate linear mixed models (mvLMM) and have been widely applied to longitudinal data analysis in animal breeding (Schaeffer, 2004). Our previous studies demonstrated the advantages of longitudinal GWAS over single trait GWAS (Ning, *et al.*, 2017). In our previous methods, we treat SNP effects as fixed regression coefficients and use a sparse matrix technique in ASReml (Gilmour, *et al.*, 2014) along with the population parameters previously determined (P3D) algorithm (Zhang, *et al.*, 2010) to reduce computing time. However, it is still computationally challenging when a marker inferred dense kinship matrix (rather than a sparse pedigree derived numerator relationship matrix) is used to capture individual genetic relationships. With the marker inferred kinship matrix, the computational complexity is *O*(*m*^3^), where *m* is the total number of phenotypic records.

To address the computational efficiency issue, we developed two efficient algorithms for longitudinal trait GWAS: fixed regression strategy with eigenvalue decomposition (Kang, et al., 2008; Lee and van der Werf, 2016; Zhou and Stephens, 2014) (GMA-fixed) and linear transformation of genomic estimation values (Gualdron Duarte, *et al.*, 2014; Ning, *et al.*, 2018) (GMA-trans) for unbalanced and balanced longitudinal traits, where unbalanced means that different individuals may be recorded at different time points and balanced means that all individuals are measured at the same time points. In order to investigate the properties of our new methods, a series of simulation studies were conducted to compare the methods with the existing univariate linear mixed model method. Furthermore, we validated our methods using an unbalanced dairy cow milk production dataset and a balanced mouse growth dataset.

## Results

### 1 Method overview

Some key features of the new methods are presented here. Details of the new methods are presented in Supplementary Note (**Additional file 1**). In the variance parameters estimation, we incorporated the expectation-maximization (EM) algorithm into the average information (AI) matrix to build a weighted information matrix (Jensen, 1997), which guarantees the variance parameters to converge rapidly within their legal domain. In the longitudinal GWAS analysis, the GMA-fixed and GMA-trans algorithms are applied in unbalanced and balanced data, respectively (Figure 1). In GMA-fixed, we treated each SNP effect as fixed regression coefficients and used the Legendre polynomials to model the time-dependent SNP effects. Similar to the studies of Kang, et al. (2010) and Zhang, et al. (2010), we estimated the variance parameters from the null model and then used these estimated parameters in subsequent analysis when markers are detected one at a time. The null model does not include the scanned SNP but it does include the polygenic effect captured with the kinship matrix. We performed eigenvalue decomposition on the phenotypic (co)variance matrix and rotated the RRM with eigenvectors. This allows us to transform the mixed model analysis into a weighted least squares analysis. The computational complexity of such a longitudinal GWAS step is reduced from *O*(*m*^3^) to *O*(*m*^2^) per-SNP. Parallel to GMA-fixed, we also performed linear transformation on the genomic estimated values in GMA-trans for unbalanced longitudinal GWAS. The basic idea in phenotype prediction is that the time varied additive genetic effect of each individual is cumulative in terms of genome-wide SNP effects. Here, we first estimated the time varied additive effects with the RRM for each individual and then transformed effects for individuals to time varied SNP effects. Wald tests were used to examine significant associations of individual SNPs with the phenotype. Compared with GMA-fixed, GMA-trans takes advantage of some intermediate results of matrix calculation in the variance parameter estimation step and avoids calculation of the phenotypic (co)variance matrix and its eigenvalue decomposition. This has reduced the computational complexity from *O*(*m*^2^) to *O*([*n*(*df* + 1)]^2^), where *n* is the number of individuals and *df* is the order of the Legendre polynomials fitting the SNP effect. To ensure convergence of the iterations in the process of variance component estimation, *df* is usually less than five and thus *n*(*df* + 1) is smaller than *m* for the usual condition of more than five measures per individual.

**Figure 1.**
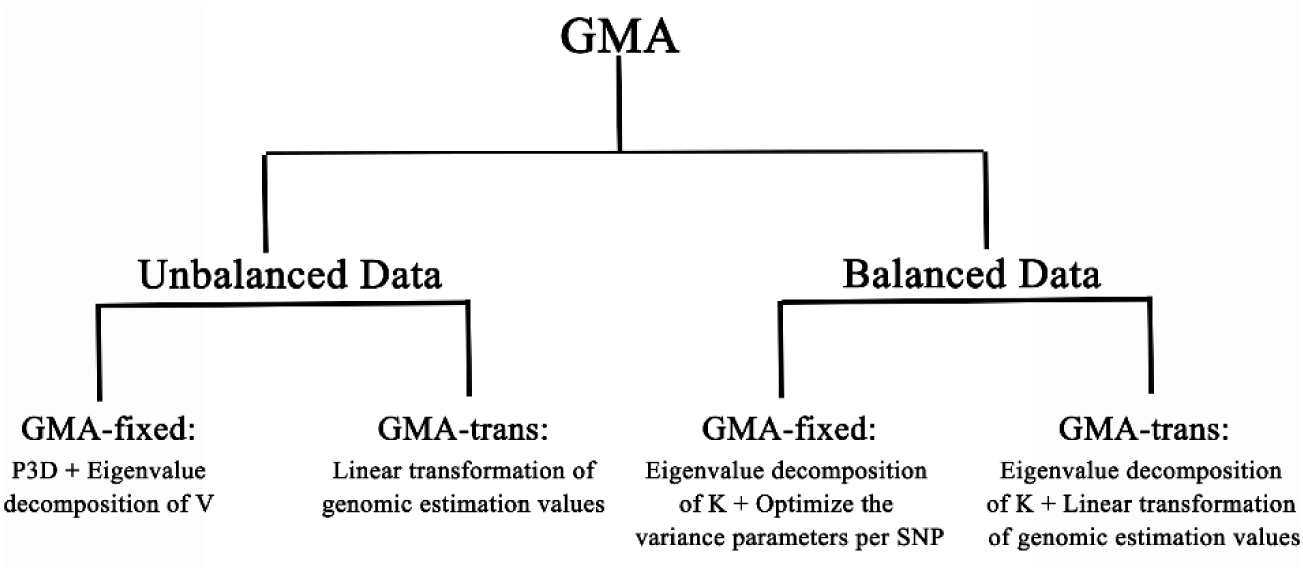
Overview of GMA for unbalanced and balanced longitudinal GWAS. P3D represents “population parameters previously determined”, which estimates the variance parameters from the null model (without SNP effects) and keeps these estimated variances as constants in the marker scanning step that follows; **V** is the phenotypic (co)variance matrix; **K** is the marker inferred relationship matrix.

Additionally, we further enhanced the GMA performance for balanced longitudinal data through eigenvalue decomposition of the genomic relatedness matrix (time complexity of O(*n*^3^)) to rotate the RRM (time complexity of O(*n*^2^)). The time complexity of variance component estimation for the rotated RRM is O(*n*) compared with O(*n*^3^) of the unbalanced longitudinal data. With the rotated RRM, we improved the QTN detection power of GMA-fixed through re-estimating the variance components for each tested SNP. The computational complexity for GMA-trans is also reduced to O(*n*) in the rotated RRM.

### 2 Simulations

We first validated the performance of GMA with simulated data. A total of four methods were compared in the simulation study. The first two methods are existing ones and the last two methods are the proposed new methods.

1. uvLMM-mean: It represents univariate linear mixed model via the mean value. Here, we analysed a random measurement each time and repeated a certain number of times for unbalanced data or analysed the measurement of each single time point separately for balanced data with the LMM method. The power estimation was obtained by taking the mean power across different analyses. We used this simulation study to obtain the empirical power of uvLMM that has ignored the time variable and thus the covariance matrix among different time points.
2. uvLMM-min: It represents univariate linear mixed model via the minimum value. The algorithm originated from Kwak, et al. (2014). With this method, we analysed a random measurement each time and repeated a certain number of times for unbalanced data or analysed one measurement for each time point separately for balanced data with the LMM method. The minimum *p*-value was used to determine the significance for a SNP.
3. GMA-trans: Linear transformation of genomic estimation values.
4. GMA-fixed: The fixed regression coefficient with eigenvalue decomposition.

To make the simulation as close as possible to reality, we perform simulations based on two real datasets, a dairy cow dataset (Ning, et al., 2017) with milk yield trait and an inter-cross F_2_ mouse dataset (Gray, *et al.*, 2015) with body weight trait. The dairy cow dataset is a large unbalanced one with 5,982 cows of 52,732 total records across days from the first lactation (from day 5 to day 305) and the total number of SNP markers is 71,527. The mouse dataset is small and balanced with 11,833 SNPs and 1,212 mice measured from week 1 to week 16 incremented by 1 week. To study the null distributions of different methods, we calculated the kinship matrix from the original SNPs and randomly shuffled each SNP across individuals when it was scanned to purposely destroy the association of the phenotypes with the scanned SNP. The *p*-values from the permuted samples are supposed to follow a uniform distribution *U*(0,1) under the null model. Figure 2 (the upper panels) shows that the type I errors are well controlled by our longitudinal GWAS algorithms and the uvLMM-mean algorithm, but are not controlled by the uvLMM-min method.

**Figure 2.**
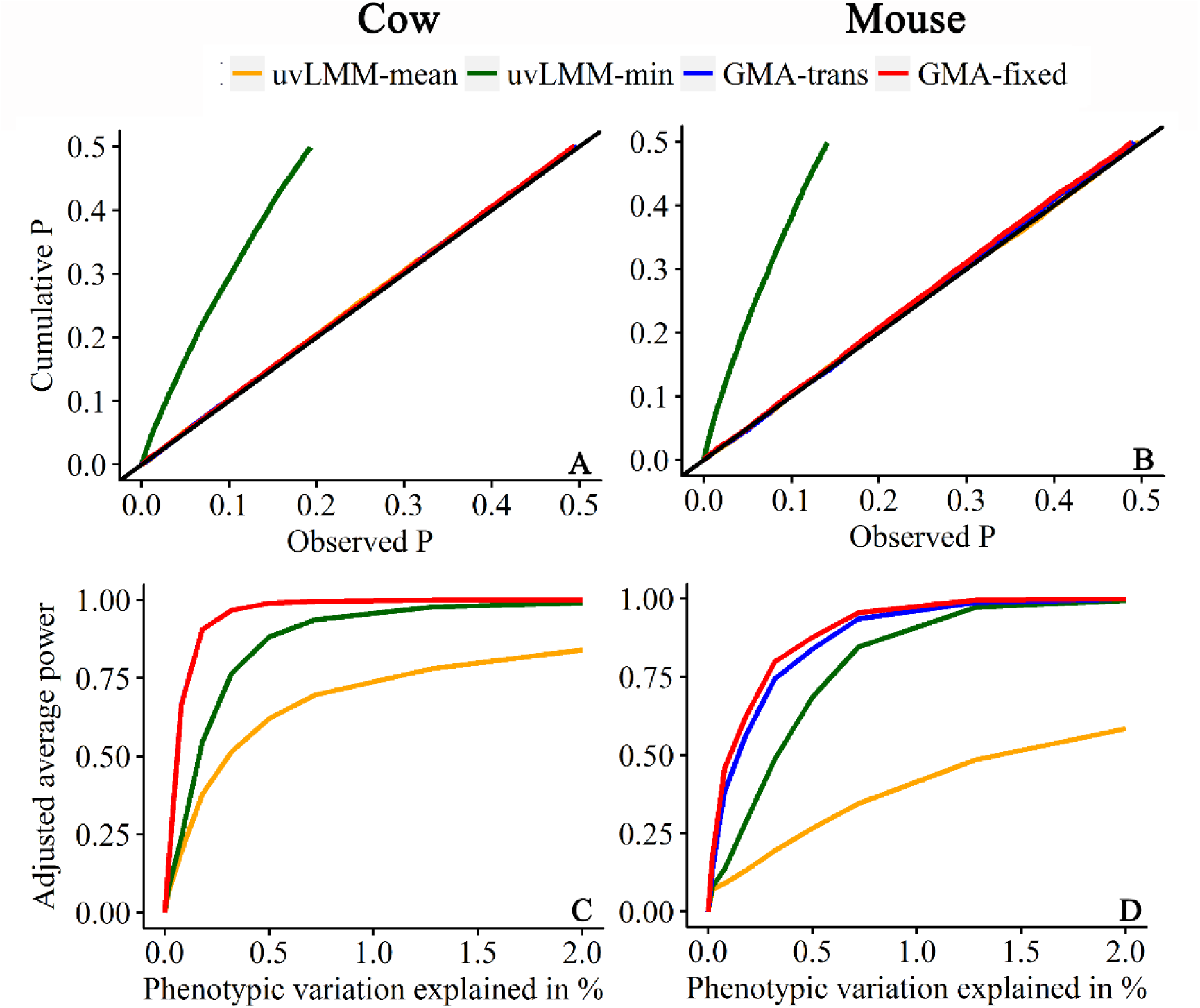
Cumulative *p*-value distributions and adjusted statistical powers of different methods in the simulation study. The left panels (A and C) represent the unbalanced dairy cow data and the right panels (B and D) represent for the balanced mouse data. The upper panels (A and B) represent distributions of the randomly shuffled SNPs. Under the null model, the cumulative *p*-value distribution should follow a uniform distribution of *U*(0,1) that overlaps with the diagonal line. Deviation from the diagonal line indicates spurious associations. The lower panels (C and D) represent the adjusted average power at different QTN contributions. The phenotypic variance is the average variance across different time points for QTN with allele frequency 0.5. The average adjusted power is calculated from 100 QTNs with nine different effects of the genetic curves. The red line overlapping with the blue line in Panel C indicates that GMA-fixed and GMA-trans have very similar power for the dairy cow data analysis.

We obtained empirical statistic powers of different methods by adding QTN effects back to the original phenotypes (Yu, et al., 2006). Nine different QTN effect functions (curves) were simulated for the unbalanced dairy cow data and the balanced mouse data (**Supplementary Figure 1** and **Supplementary Figure 2**). The results are illustrated in Figure 2 (the lower panels) showing that the new methods have higher power than two uvLMM methods. In particular, the approximate GMA-fixed algorithm for the unbalanced data has almost the same power as GMA-trans, while the exact GMA-fixed algorithm for the balanced data (optimize variance parameters for each SNP) has the highest power. The uvLMM-mean algorithm has the lowest statistic power, which demonstrates the benefit of using the new GWAS methods of longitudinal traits.

### 3 Application to real data

Prior to scanning markers in the GWAS, we first compared our efficient algorithms for variance component estimation to two existing methods, Wombat (Meyer, 2007) and MTG2 (Lee and van der Werf, 2016) (Table 1). In variance component estimation, the Wombat program uses a hybrid algorithm consisting of a few initial rounds of PX-EM (Liu, *et al.*, 1998), followed by the AI algorithm, while MTG2 uses the pure AI algorithm with eigenvalue decomposition technique and moderates the magnitude of updates when the parameters go outside the legal domain of the parameter space. In general, the GMA methods converged faster with fewer iterations than the two methods. For the balanced longitudinal mouse data, our algorithm took only 2 seconds to complete the analysis while MTG2 took 5 seconds and Wombat took 40 minutes. Even for unbalanced longitudinal dairy cow data, the GMA method was substantially faster than Wombat.

**Table 1.** Computational times of different methods for variance component estimation (including iteration number) and the subsequent step of GWAS.

| Category | Method | Computational time |  |
| --- | --- | --- | --- |
|  |  | Mouse data | Dairy cow data |
| Variance estimation | Wombat | 40 min (11) | 105 (12) |
|  | MTG2 | 5 s (15) | - |
|  | GMA | 2 s (9) | 5.3 h (7) |
| GWAS | uvLMM | 14.4 min | 3.7 h |
|  | GMA-fixed | 5.1 h | 16.5 h |
|  | GMA-trans | 1.7 min | 3.8 h |
|  | GMA-trans + GMA-fixed | 7 min | - |

All computations were performed on Intel Xeon E5 2.2 GHz CPU. We used the third order of Legendre polynomials for the mouse dataset and the forth order for dairy cow dataset. The same convergence criterion was used for all methods in variance estimation, where the iteration stopped when the difference of the log likelihood values between consecutive iterations is smaller than 0.001. The uvLMM method was implemented in the GEMMA (Zhou and Stephens, 2012) package. In variance component estimation, the Wombat program uses a hybrid algorithm consisting of a few initial rounds of PX-EM(Liu, et al., 1998), followed by the AI algorithm; MTG2 uses the pure AI algorithm and moderates the magnitude of updates when the parameters go outside the legal domain of the parameter space; GMA incorporates the EM algorithm into the AI matrix to build a weighted information matrix.

We now compared results of the longitudinal GWAS obtained via the GMA-trans and uvLMM method. The two took about the same amount of time for the unbalanced data, but GMA-trans is much faster than uvLMM for the balanced data. Furthermore, the current GMA-trans algorithm for unbalanced data is several times faster than the GMA-fixed algorithm. We compared the *p*-values from GMA-fixed and GMA-trans and discovered that they are exactly the same (**Supplementary Figure 3, Panel A**). For the balanced mouse data, GMA-fixed optimizes the variance components per SNP and is much slower than GMA-trans. However, the correlation coefficient of the *p*-values between the two methods is very high (Pearson’s *r* = 0.995). The *p*-values of GMA-fixed are often smaller than the *p*-values of GMA-trans (**Supplementary Figure S3, Panel B**), which means that GMA-fixed may detect more loci than GMA-trans. Taking into account the fast computational speed of GMA-trans and the high power of GMA-fixed (due to re-estimation of variance components), we pre-selected SNPs based on a relaxed *p*-value criterion, say *p*-value < 0.01, from GMA-trans and then recalculated the *p*-values from GMA-fixed. As a result, the lost power by GMA-trans has been be rescued by GMA-fixed (**Supplementary Figure S3, Panel C**), yet the reduced computational time remained at the same level (about 7 minutes) as the GMA-trans method.

For the unbalanced dairy cow data, both GMA-fixed and GMA-trans identified four significant SNPs (three at 1.65-1.81Mb and one at about 4.36Mb on chromosome 14) for milk yield without inflated false positives after multiple test correction using false discovery rate (FDR) with FDR < 5% (*q* value < 0.05) (**Supplementary Figure 4**). One of the SNPs (1,801,116bp) is located within the *DGAT1* gene (1,795,351-1,804,562bp) that is reported to be a major gene affecting milk production traits (Grisart, *et al.*, 2004), and all significant SNPs are within the boundary of the reported QTL for milk yield (Hu, *et al.*, 2015). We compared the additive effect curves of the four significant SNPs with milk yield trajectory in **Supplementary Figure 5** and found very similar patterns between the curves, though the peak time of SNP effects (at about 200 days) is delayed compared to the peak time of the phenotypic trajectory (at about 80 days). The results indicate that *DGAT1* exhibits its main effects after the lactation peak and may contribute to the persistency of milk production (Strucken, *et al.*, 2015).

For the balanced mouse data, GMA-fixed detected two candidate regions (112-128Mb on chromosome 10 and 75-88Mb on chromosome 13; *q* value < 0.05) (Figure 3A,B), while GMA-trans only detected one of the two regions (119-125Mb on chromosome 10; *q* value < 0.05) (Figure 3C,D). In this study, we also used the uvMLM-min method for comparison. The quantile-quantile (Q-Q) plot in Figure 3E shows that uvMLM-min appears to have higher type I errors than GMA, which is consistent with the simulation study. We then used the permutation test to determine the *p-*value threshold (genome-wide significance level of 0.05) for declaration of significance. This criterion led to the detection of one candidate region (118-125Mb on chromosome 10) (Figure 3F). Meanwhile, we compared the additive effect curves of the significant SNPs with the phenotypic trajectory (Figure 4). The additive effect curves of significant SNPs on chromosome 10 have patterns similar to the phenotypic trajectory. The region has also been reported as a candidate QTL by Gray, et al. (2015). However, the additive effect curves of the new candidate QTL on chromosome 13 are concave in shape and the QTL effect is inverse in the interim compared to the beginning and end (Figure 4C).

**Figure 3.**
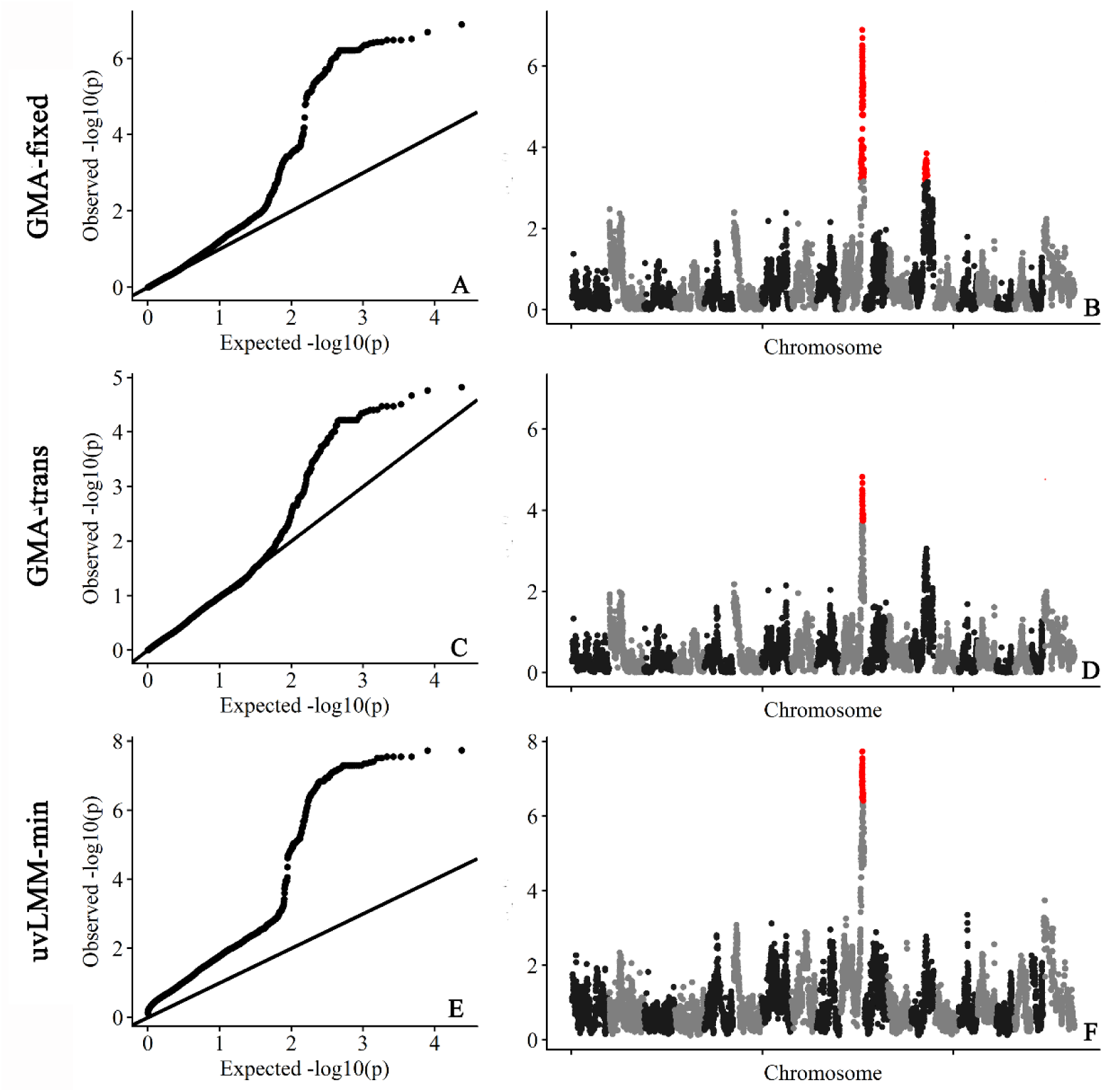
Association studies of growth trajectory in the mouse population with the GMA-fixed method (panels at the top), the GMA-trans method (panels in the middle) and the uvMLM-min method (panels at the bottom).

**Figure 4.**
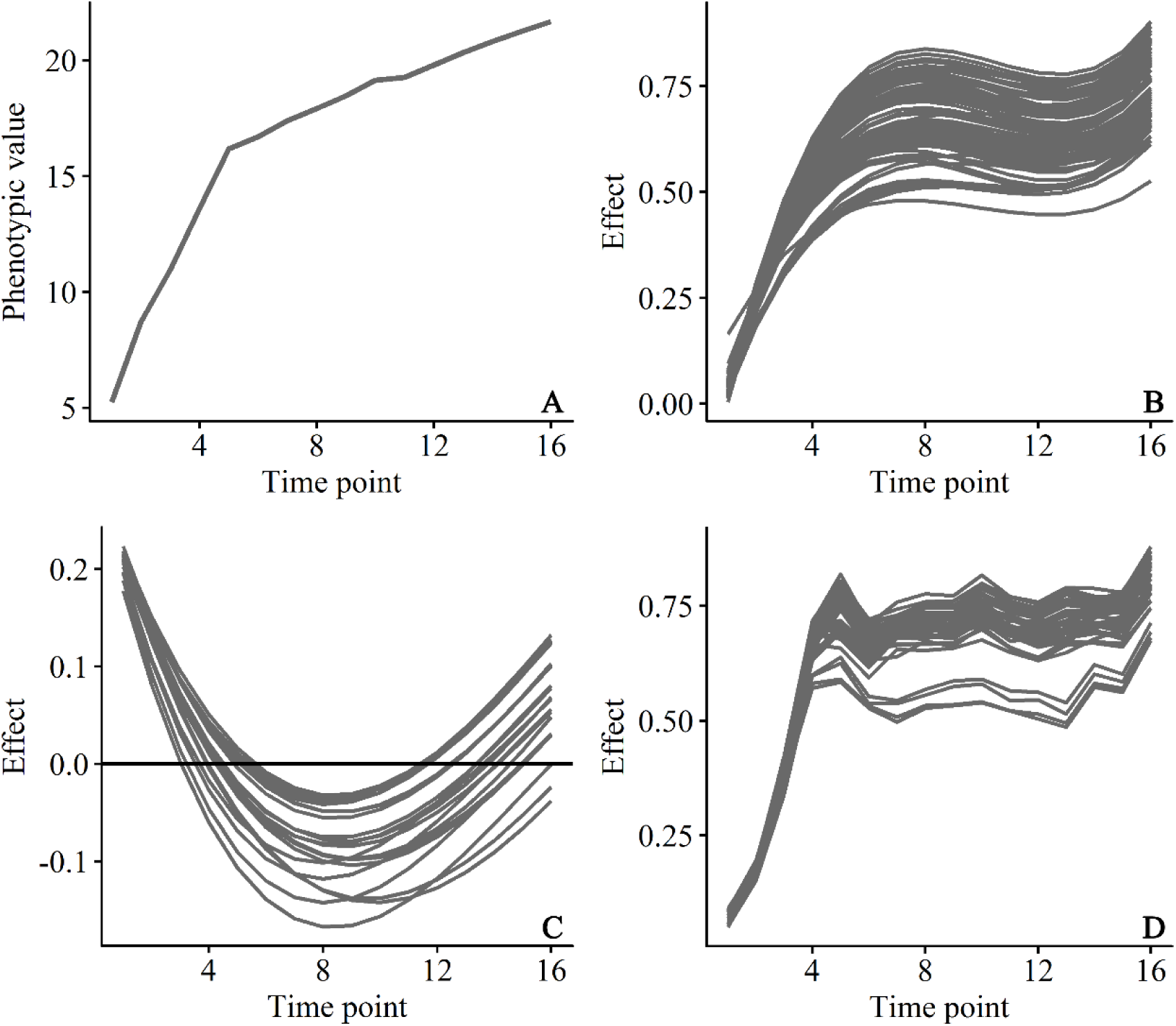
The phenotypic and significant SNPs changing pattern for body weight in the mouse data. (A) The average phenotypic values plotted against age (from week 1 to week 16 incremented by 1); (B) The predicted growth trajectories of QTL effects for all significant SNPs between 112Mb and 128Mb on chromosome 10 by the GMA-fixed method; (C) The predicted growth trajectories of QTL effects for all significant SNPs between 75Mb and 88Mb on chromosome 13 by the GMA-fixed method; (D) The predicted growth trajectories of QTL effects for all significant SNPs between 118Mb and 125Mb on chromosome 10 by uvMLM-min method.

## Discussion

Longitudinal GWAS provides an appealing approach to probe the dynamic genetic mechanism of complex traits. However, successful application of the longitudinal GWAS is challenged by cryptic genetic relationship, dependency among the time course observations and time-consuming computation challenge. Here, we developed efficient analysis algorithms for longitudinal GWAS dealing with either balanced or unbalanced longitudinal data. Our algorithms are based on RRM, a multivariate linear mixed model (mvMLM). The RRM includes a time varied polygenic effect and a permanent environmental effect to explain the cryptic genetic relationship and dependency among observations. To improve the computational efficiency, we built a weighted information matrix from the EM algorithm and the AI information matrix, which guarantee the variance parameters to converge with fewer iterations. In the meantime, we proposed the fixed regression coefficient approach accompanied with eigenvalue decomposition strategy (GMA-fixed) and linear transformation of genomic estimation values (GMA-trans) algorithms. Simulations based on genotypes and phenotypes of actual populations show that our algorithms perform very well in terms of high statistical power and low false positive rate compared with the conventional uvLMM implemented GWAS. Application to the unbalanced dairy cow data and the balanced mouse data further validated the benefits of our longitudinal GMA.

There are various dynamic patterns of genetic controls represented by permanent QTLs, early QTLs, late QTLs and inverse QTLs (Wu and Lin, 2006). In this study, we used Legendre polynomials to model the dynamic changing process of QTL. This is a non-parametric approach because it makes no assumption about the shape of the curve. The method also reduces the correlations between the estimated random regression coefficients so that variance parameter estimation converges very rapidly. From the analyses of the two real data, we observed that the main QTLs tend to have similar changing patterns with the phenotypic curve, indicating that these QTLs determine the dynamic genetic mechanism of longitudinal traits. We also identified an inverse QTL (one genotype performs better than the other during early stage of growth, but the other genotype performs better during later stage of the growth) for the mouse data with GMA-fixed. These QTLs and others with minor effects can play a regulation role in shaping the final phenotypic trajectory.

For balanced data, GMA-fixed is more powerful than GMA-trans because it optimizes the variance parameters per SNP, but the latter is much faster. The GMA-trans step followed by the GMA-fixed step is recommended because it takes advantage of the high power of GMA-fixed and the high speed of GMA-trans. For unbalanced data, it is time consuming to optimize the variance components for each SNP. Since GMA-fixed and GMA-trans have similar power for unbalanced data, GMA-trans is recommended.

In contrast to uvLMM with only two variance parameters (additive and residual variances), RRM has a complicated covariance structure with many variance parameters (depending on the orders of the Legendre polynomials). As a result, RRM may need more iterations to converge and, sometime, may encounter a convergence issue. If the iteration process stops early before convergence, the GMA algorithms may be subject to a higher Type I error. The orders of the Legendre polynomials can be determined by a model selection criteria, such as Akaike information criterion (Akaike, 1974) (AIC) and Bayesian information criterion (Schwarz, 1978) (BIC). To avoid any convergence issue, three or four orders of Legendre polynomials are recommended in practice. If the GMA algorithm encounters convergence issue even with low order of Legendre polynomials, the GMA-trans algorithm with an increased iteration number in variance parameter estimation step is recommended.

In our study, we focus on the traits changing over time. However, our developed GMA algorithm can be naturally applied to traits changing with other dynamic environmental covariates, such as solar radiation, solar radiation and temperature. Modern automatic information platforms can record abundant environmental data, while advanced genotyping technologies allow accessing to genomic information on a large scale. The GMA can utilize the two types of high dimensional information to tackle genome-wide genotypes and environments (G×E) interactions efficiently, which facilitates dissecting the complex genetic architecture of dynamic traits.

## Methods

### 1 Data

Two datasets were analysed in the study: a mouse data (Gray, et al., 2015) and a dairy cow data (Ning, et al., 2017). The mouse data contain 1,212 F_2_ from the cross between the Gough Island mice and the WSB/EiJ strain. The body weight trait was measured from week 1 to week 16 incremented by 1 week (16 measurements per mouse). There are 11,833 available SNP markers across the mouse genome after proper quality control. The dairy cow data include 5,982 individual cows. The milk yield trait of the first parity were analysed in this study. The cows with less than six records were filtered out, which resulted a total of 52,732 records. The SNPs with a minor allele frequency (MAF) less than 0.03 and a failed the Hardy-Weinberg equilibrium (HWE) test (*p*-value < 10^−6^) were removed, resulting in 71,527 SNPs for the subsequent longitudinal GWAS analyses.

### 2 Simulation

In order to assess the null distributions of different models, we calculated the kinship matrix from the original SNPs and randomly shuffled each SNP across individuals when it was scanned to purposely destroy the association of the phenotypes with the scanned SNP. The covariance structure of original phenotypes induced by the complex cryptic genetic relationship among the individuals will not be disorganized in this way. Under the expectation that random SNPs are unlinked to polymorphisms controlling these traits, the cumulative *p*-value distribution follows a uniform distribution of *U*(0, 1). The empirical power was obtained from populations simulated from the genotypes of the current populations (the mouse and the cattle data) by assigning genetic effects to selected markers and adding maker effects back to the original phenotypes (Yu, et al., 2006), *i.e.*, *y*_*i*,*new*_(*t*) = *y*_*i*_(*t*) + *s*_*i*_*SNP*(*t*). Where *y*_*i*_(*t*) is the observed phenotypic value of individual *i* at time *t*; *s*_*i*_ is a genotype indicator for individual *i* which is assigned 0, 1 and 2 for genotype *aa*, *Aa* and *AA*, respectively; *SNP*(*t*) represents the simulated time varied effect for selected marker; *y*_*i*,*new*_(*t*) is the newly generated phenotypic value of individual *i* at time *t*. We random selected 100 SNPs from the genome and assigned then with nine different maker effect curves. The time varied SNP effects were then adjusted so that they contributed to some predetermined proportions of the phenotypic variance (average proportion across the time points, 0.02-2% at MAF of 0.5). The genetic effect curves were assigned to the 100 random selected SNPs, one at a time. The simulated data were analysed by the proposed new methods and existing methods. A marker was declared as significant if the *p*-value was smaller than the empirical threshold (the 5^th^ percentile of the null distribution).

### 3 GMA algorithms

Details of the GMA algorithms are described in Supplementary Note (**Additional file 1**).

## Acknowledgements

The project was supported by the National Natural Science Foundations of China (31661143013), Changjiang Scholars and Innovative Research Team in University (IRT_15R62) and Jinxinnong Animal Science Development Foundation. The authors are grateful to Jian Yang for his comments on an early version of the manuscript.

## Author contributions

C.N. and J.F.L. conceived and designed the experiments. C.N. and D.W. contributed analytic tools and analysed the data. L.Z., J.W., H.K., S.Z, X.Z. and S.X. participated in the result interpretation and paper revision. C.N. and J.F.L. wrote the paper with comments from X.Z. and S.X. All authors read and approved the final manuscript.

## Competing interests

The authors declare that they have no competing interests.

## Additional file

Additional file 1: Supplementary Figure 1-5 and Supplementary Note.

## Reference

Akaike, H. (1974) A new look at the statistical model identification, IEEE transactions on automatic control, 19, 716–723.

Fahlgren, N., Gehan, M.A. and Baxter, I. (2015) Lights, camera, action: high-throughput plant phenotyping is ready for a close-up, Current opinion in plant biology, 24, 93–99.

Gilmour, A., et al. (2014) ASReml user guide. Release 4.1 structural specification, VSN International Ltd, Hemel Hempstead, HP1 1ES, UK www.vsni.co.uk.

Gong, Y. and Zou, F. (2012) Varying coefficient models for mapping quantitative trait loci using recombinant inbred intercrosses, Genetics, 190, 475–486.

Gray, M.M., et al. (2015) Genetics of Rapid and Extreme Size Evolution in Island Mice, Genetics, 201, 213–228.

Grisart, B., et al. (2004) Genetic and functional confirmation of the causality of the DGAT1 K232A quantitative trait nucleotide in affecting milk yield and composition, Proceedings of the National Academy of Sciences, 101, 2398–2403.

Gualdron Duarte, J.L., et al. (2014) Rapid screening for phenotype-genotype associations by linear transformations of genomic evaluations, BMC bioinformatics, 15, 246.

Hu, Z.-L., Park, C.A. and Reecy, J.M. (2015) Developmental progress and current status of the Animal QTLdb, Nucleic acids research, gkv1233.

Jensen, J. (1997) Residual maximum likelihood estimation of (co) variance components in multivariate mixed linear models using average information, Journal of the Indian Society of Agricultural Statistics, 49, 215–236.

Kang, H.M., et al. (2010) Variance component model to account for sample structure in genome-wide association studies, Nature genetics, 42, 348–354.

Kang, H.M., et al. (2008) Efficient control of population structure in model organism association mapping. Genetics. pp. 1709–1723.

Kellogg, E.C., Thrasher, A. and Yoshinaga-Itano, C. (2014) Early predictors of autism in young children who are deaf or hard of hearing: three longitudinal case studies, Seminars in speech and language, 35, 276–287.

Kwak, I.Y., et al. (2014) A simple regression-based method to map quantitative trait loci underlying function-valued phenotypes, Genetics, 197, 1409–1416.

Lee, S.H. and van der Werf, J.H. (2016) MTG2: an efficient algorithm for multivariate linear mixed model analysis based on genomic information, Bioinformatics, 32, 1420–1422.

Li, Z. and Sillanpaa, M.J. (2015) Dynamic Quantitative Trait Locus Analysis of Plant Phenomic Data, Trends in plant science, 20, 822–833.

Lippert, C., et al. (2011) FaST linear mixed models for genome-wide association studies, Nature methods, 8, 833–835.

Liu, C., Rubin, D.B. and Wu, Y.N. (1998) Parameter expansion to accelerate EM: The PX-EM algorithm, Biometrika, 85, 755–770.

McSweeney, J., et al. (2014) Predicting coronary heart disease events in women: a longitudinal cohort study, The Journal of cardiovascular nursing, 29, 482–492.

Meyer, K. (2007) WOMBAT: a tool for mixed model analyses in quantitative genetics by restricted maximum likelihood (REML), Journal of Zhejiang University. Science. B, 8, 815–821.

Ning, C., et al. (2017) Performance Gains in Genome-Wide Association Studies for Longitudinal Traits via Modeling Time-varied effects, Scientific reports, 7.

Ning, C., et al. (2018) A rapid epistatic mixed-model association analysis by linear retransformations of genomic estimated values, Bioinformatics, 34, 1817–1825.

Porto, S.M.C., et al. (2015) The automatic detection of dairy cow feeding and standing behaviours in free-stall barns by a computer vision-based system, Biosystems Engineering, 133, 46–55.

Schaeffer, L.R. (2004) Application of random regression models in animal breeding, Livest Prod Sci, 86, 35–45.

Schwarz, G. (1978) Estimating the dimension of a model, The annals of statistics, 6, 461–464.

Strucken, E.M., Laurenson, Y.C. and Brockmann, G.A. (2015) Go with the flow-biology and genetics of the lactation cycle, Front Genet, 6, 118.

Wu, R. and Lin, M. (2006) Functional mapping—how to map and study the genetic architecture of dynamic complex traits, Nature Reviews Genetics, 7, 229–237.

Xiong, H., et al. (2011) A flexible estimating equations approach for mapping function-valued traits, Genetics, 189, 305–316.

Yang, J., et al. (2011) GCTA: a tool for genome-wide complex trait analysis, Am J Hum Genet, 88, 76–82.

Yu, J., et al. (2006) A unified mixed-model method for association mapping that accounts for multiple levels of relatedness, Nature genetics, 38, 203–208.

Zhang, Z., et al. (2010) Mixed linear model approach adapted for genome-wide association studies, Nature genetics, 42, 355–360.

Zhou, X. and Stephens, M. (2012) Genome-wide efficient mixed-model analysis for association studies, Nature genetics, 44, 821–824.

Zhou, X. and Stephens, M. (2014) Efficient multivariate linear mixed model algorithms for genome-wide association studies, Nature methods, 11, 407–409.

