## Additional file 1 for "Efficient Multivariate Analysis Algorithms for Longitudinal Genome-wide Association Studies"

**Supplementary Information**

1. **Supplementary Figures**

**Supplementary Figure 1**

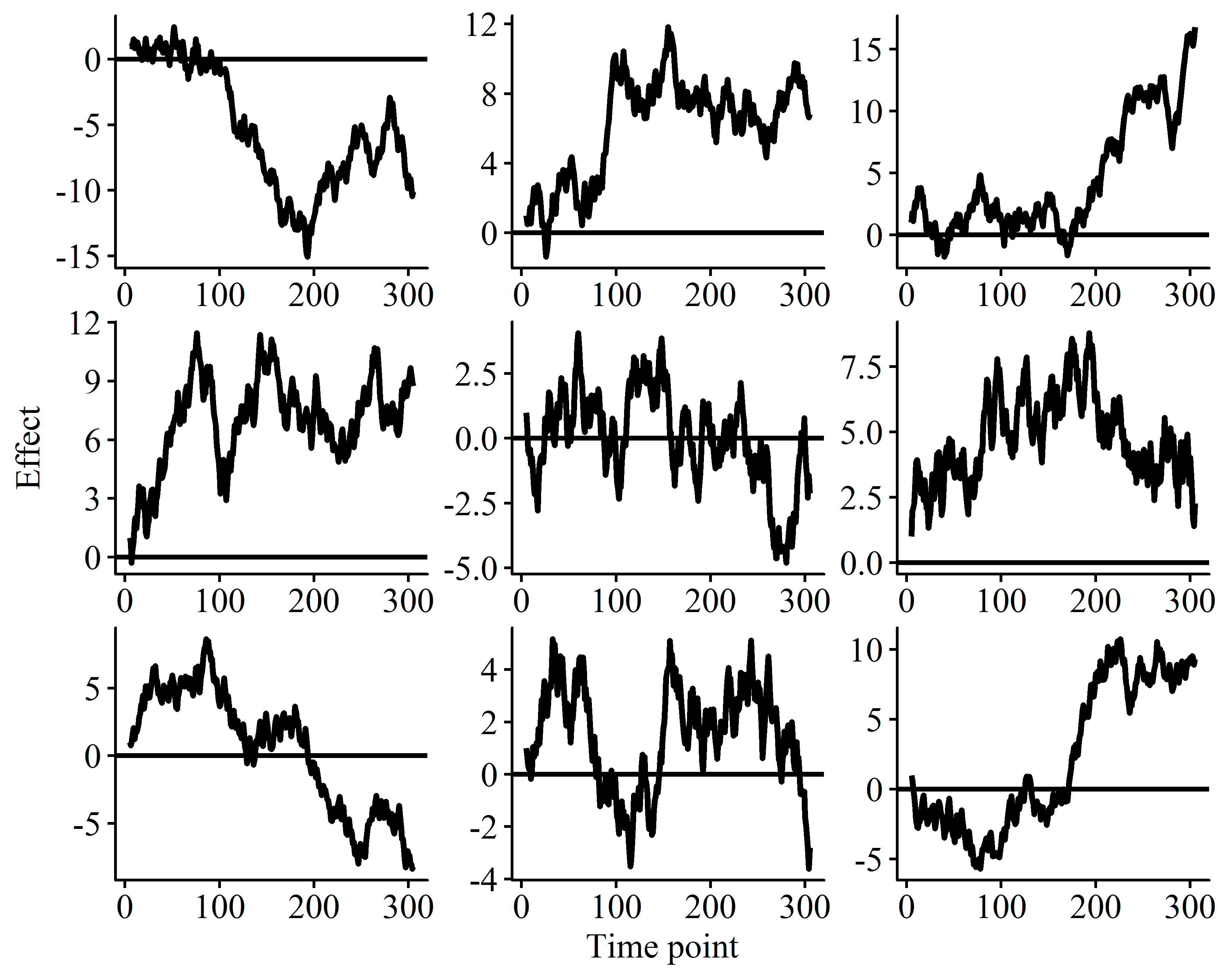

**Supplementary Figure 1 Nine different simulated marker effect curve in the dairy cow data.**

**Supplementary Figure 2**

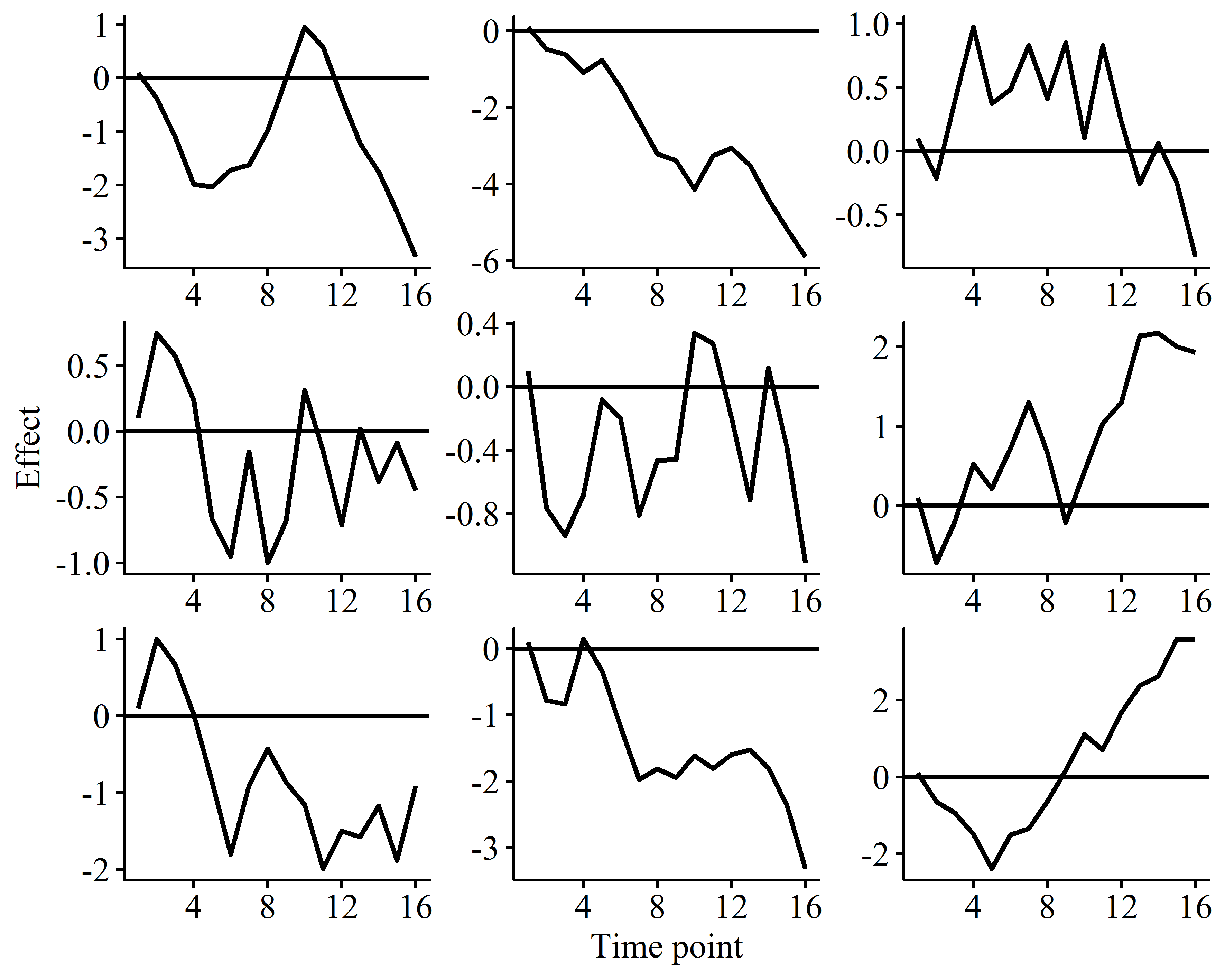

**Supplementary Figure 2 Nine different simulated marker effect curve in the mouse data.**

**Supplementary Figure 3**

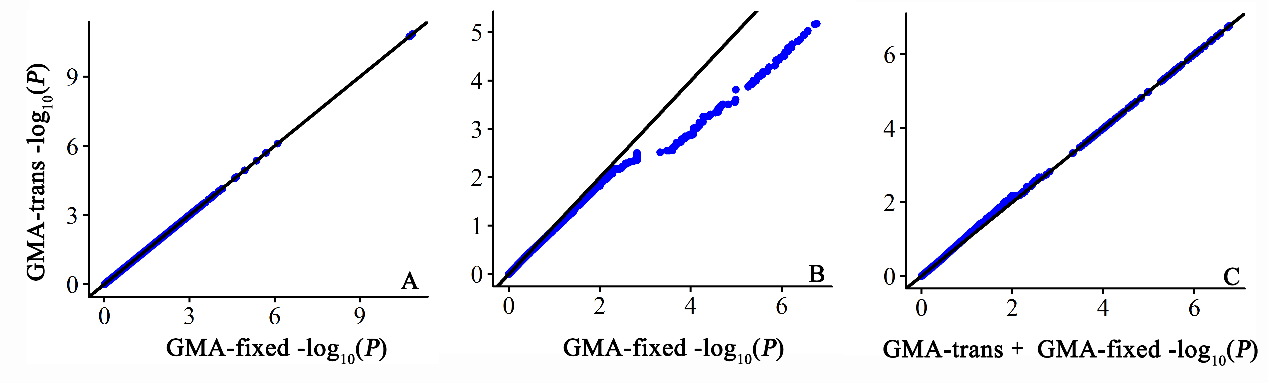

**Supplementary Figure 3 The comparison of *P*-values (in -log10 scale) between GMA-fixed and GMA-trans in dairy cow data (a), GMA-fixed and GMA-trans in mouse data (b), as well as GMA-trans + GMA-fixed and GMA-trans in mouse data (c).**

**Supplementary Figure 4**

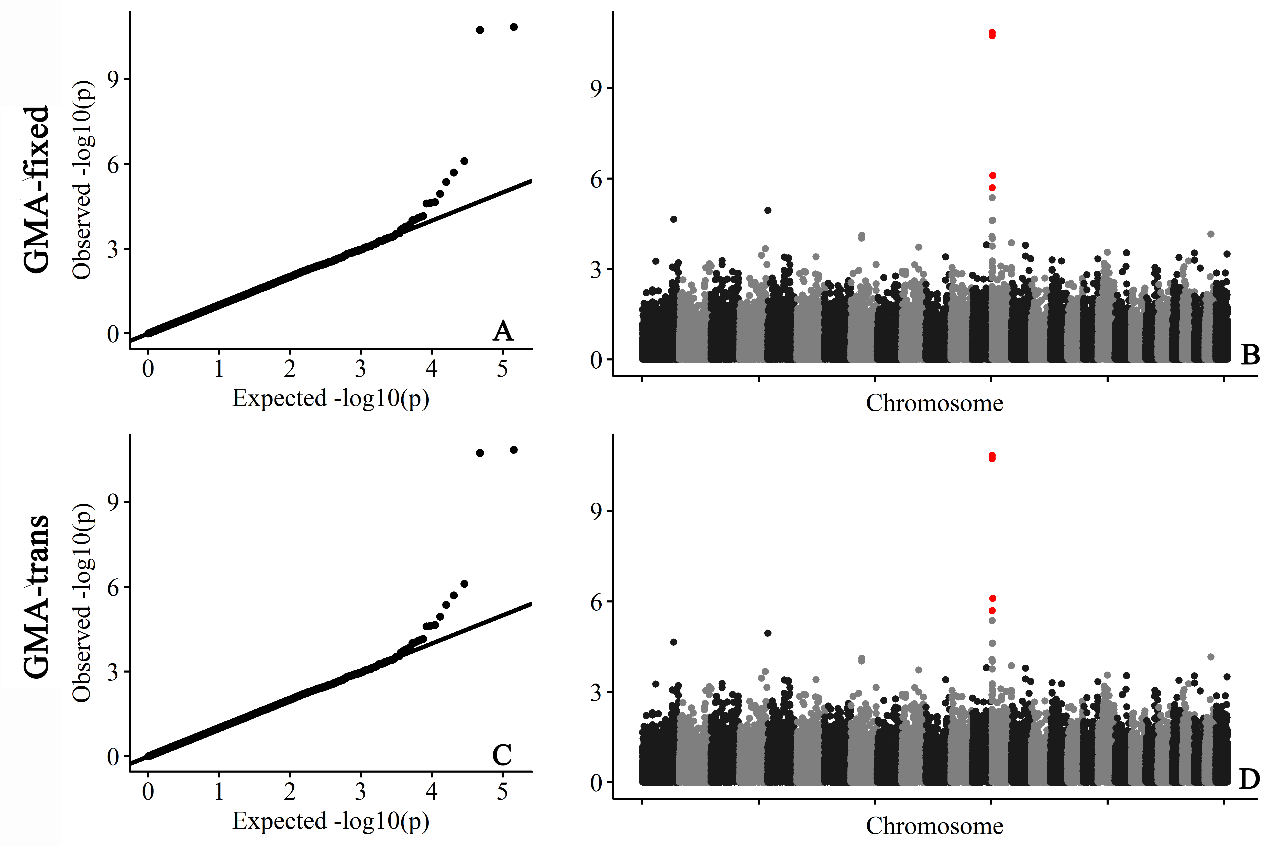

**Supplementary Figure 4 Association studies of milk yield in the dairy cow data with GMA-fixed (top) and GMA-trans (bottom).**

**Supplementary Figure 5**

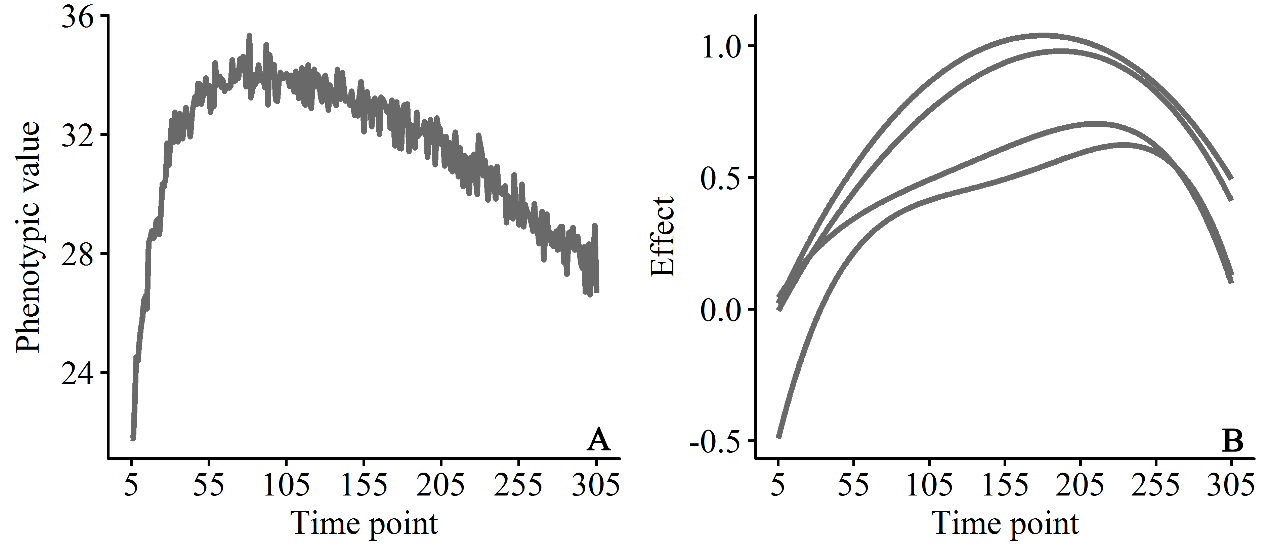

**Supplementary Figure 5 The** **phenotypic and significant SNPs changing pattern for milk yield in the dairy cow data.** (A) The average phenotypic values during the first lactation period (from day 5 to 305); (B) The predicted changing pattern for the four significant SNPs on chromosome 14 by GMA-fix method.

2. **Supplementary Note**
   2.1 **Unbalanced Longitudinal Data**
       2.1.1 **Random Regression Model**

Following Mrode (2014), a typical random regression model (RRM) can be written as

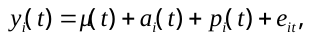
 (1)

where
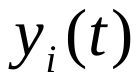
 is the phenotypic value of individual *i* at time *t*;
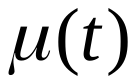
 is the overall mean at time *t*;
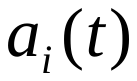
 and
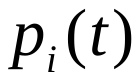
 are the time-varied additive genetic effect and permanent environmental effect respectively for individual *i*;
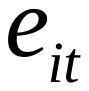
 is the time-independent random residual for each measurement of individual *i* at time *t*. Here,
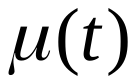
,
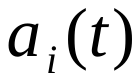
 and
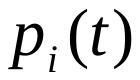
 can be denoted as the linear regression for a set of basis functions, such as Legendre polynomials,

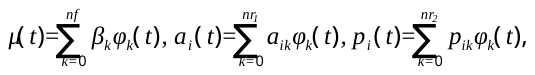
 (2)

where *nf*,
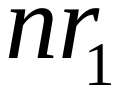
 and
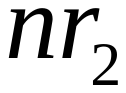
 are the orders of corresponding basis functions;
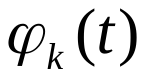
 is the value of the *k*th basis function at time *t*;
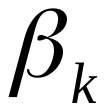
 is the *k*th fixed regression coefficient;
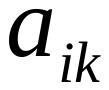
 and
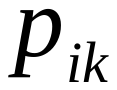
 are the *k*th random regression coefficients for additive genetic effect and permanent environmental effect of the *i*th individual. The matrix form for the *i*th individual can be represented as

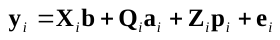
 (3)

Here, we assume there are *n* individuals and the number of records for individual *i* is
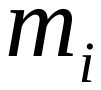
 (
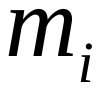
 is different for each individual), then the total number of records for all individuals is
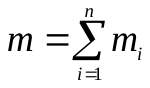
. Thus,
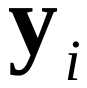
is a
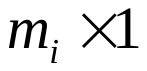
 vector of phenotypic values for individual *i*; **b** is a
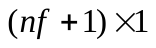
 vector of fixed regression coefficients;
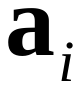
 is a
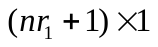
 vector of random regression coefficients for additive genetic effects of individual *i*;
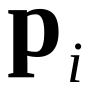
 is a
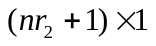
 vector of random regression coefficients for permanent environmental effects of individual *i*;
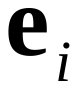
 is a
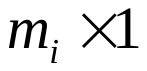
 vector of random residuals.
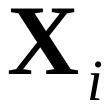
,
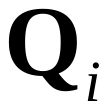
 and
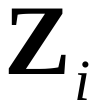
 are the corresponding design matrices. The corresponding vectors and matrices are formulated as follows

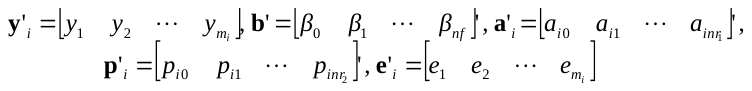
 (4)

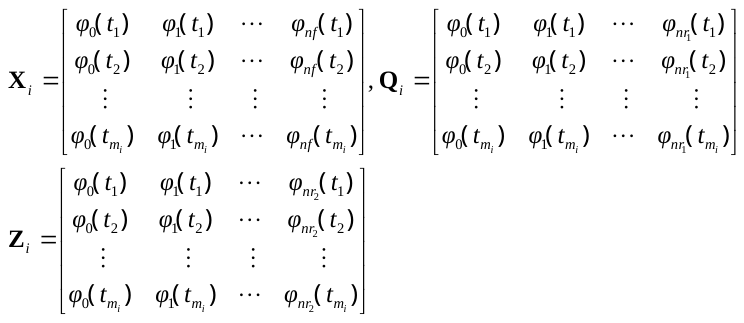
 (5)

The (co)variance matrices of all random effects is

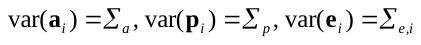
 (6)

Here,
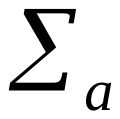
 is the (co)variance matrix for random regression coefficients of additive polygenic effects with size of
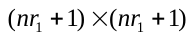
;
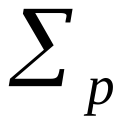
 is the variance–covariance matrix of random regression coefficients for permanent environmental effects with size of
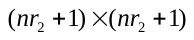
;
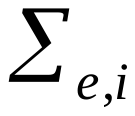
 is a diagonal matrix with different values at different time periods.

The matrix form for the all individuals can be represented as

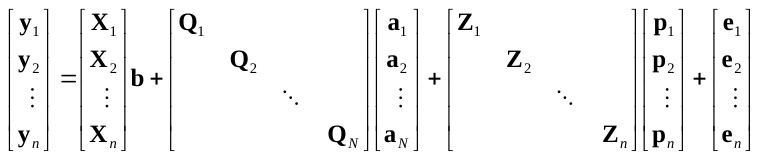
 (7)

Here, we further abbreviate the matrix as

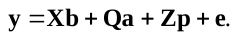
 (8)

The (co)variance matrices of all random effects is

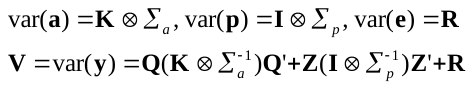
 (9)

Here, **K** is a *N* × *N* marker-derived relationship matrix, where *N* is the number of individuals with SNP information (*N*≥*n*); **I** is the *n* × *n* identity matrix;
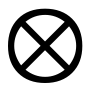
 is the Kronecker product. Matrix **K** was built with the method of VanRaden (2008) as follows

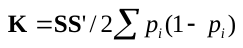
 (10)

Where *p_i_* is the second allele (allele *a*) frequency at locus *i*, and **S** is a centralized maker matrix with the *i*th column formulated as

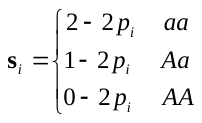
 (11)

The mixed model equations (MME) can be expressed as

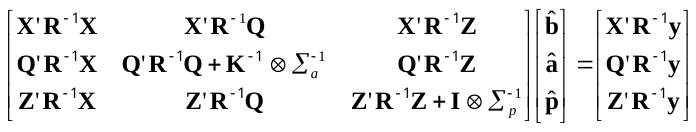
 (12)

Here, we define matrix **C** as the left hand coefficient matrix of Equation (12) and its inversion as

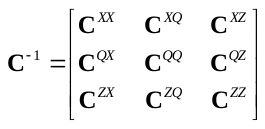
 (13)

2.1.2 **Restricted Log-likelihood Function**

According the study of Verbyla (1990), the restricted log-likelihood function can be formulated as

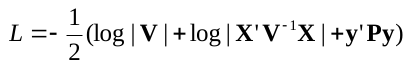
 (14)

Where,
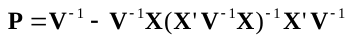
 (15)

This is called direct REML by Lee and van der Werf (2006). An alternative formula based on the mixed model equations (MME) is formulated as

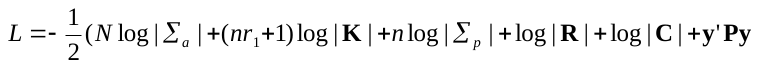
 (16)

2.1.3 **AI** **Algorithm**

Gilmour*, et al.* (1995) and Johnson and Thompson (1995) introduced an efficient average information (AI) algorithm to maximize restricted log-likelihood function and estimate variance components, which is formulated as

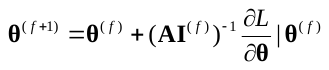
 (17)

Here, **θ** is a vector of variance components including the unique values in
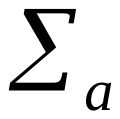
,
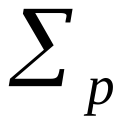
 and **R**; *f* is the iteration round;
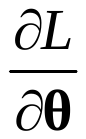
 is a vector of the first derivatives of the log likelihood function with respect to each variance component; **AI** is the average information matrix.

The first derivative for each variance covariance component *i* is

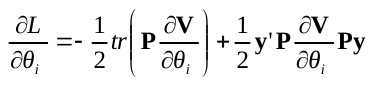
 (18)

The element of AI matrix for variance covariance component *i* and *j* is

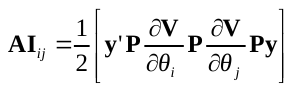
 (19)

If we define working variables for
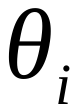
 as

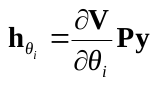
 (20)

and the matrix
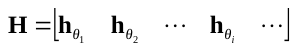

Then the AI matrix can be rewritten as

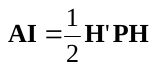
 (21)

Similar to restricted log-likelihood function, alternative formulas based on the mixed model equations for first derivatives are

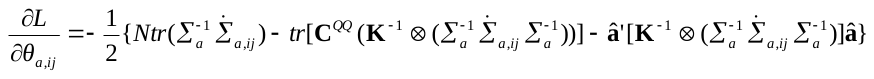
(22)

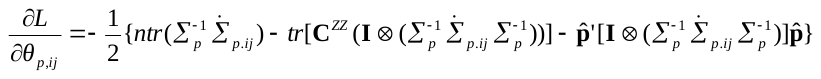
 (23)

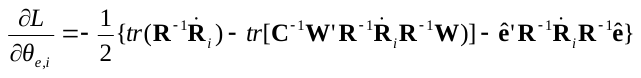
 (24)

Here, we denote
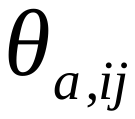
 as the *ij*th element of
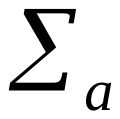
,
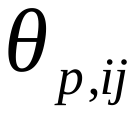
 as the *ij*th element of
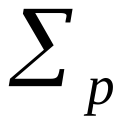
,
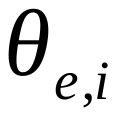
 as the *i*th unique element of **R**. Furthermore,

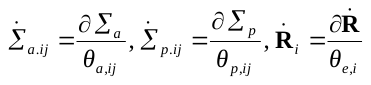
. (25)

Similarly, alternative forms for working variables is

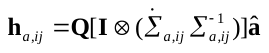
 (26)

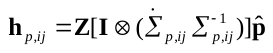
 (27)

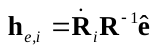
 (28)

2.1.4 **EM** **Algorithm**

Following the study of Jensen*, et al.* (1997), EM updates which are similar to the AI updates (Equation 17) can be expressed as

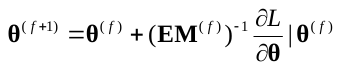
 (29)

Here, the inversion of matrix **EM** is a block diagonal matrix and the elements corresponding to
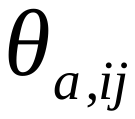
 and
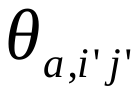
,
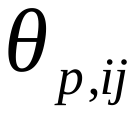
 and
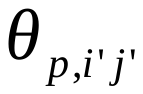
, as well as
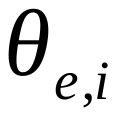
 and
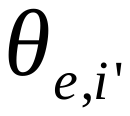
 are

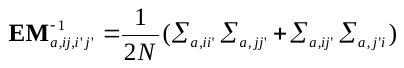
 (30)

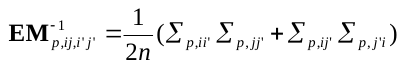
 (31)

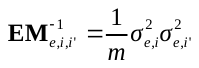
 (32)

2.1.5 **Combined EM and AI** **Algorithm**

The AI algorithm is computationally efficient. However, the estimates can go outside parameter space with illegal or bad starting values. One solution is using EM algorithm in initial iterations to obtain a better set of starting values. Unfortunately, the increase in likelihood can be very small and it takes a long time to switch to AI algorithm. Jensen, et al. (1997) provided a better alternative which use a combined information matrix

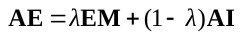
 (33)

Where λ∈[0, 1]. If λ= 0, it is a pure AI algorithm, and if λ= 1, it is a pure EM algorithm. We try λ value starting from 0 with a step (such as 0.1) until estimated variance components are within parameter space for each iteration.

2.1.6 **Variance Components Estimation for Unbalanced Longitudinal Data**

In our study, we estimate variance components using MME-based REML method for unbalanced longitudinal data. In other words, we estimate the value of restricted log-likelihood function with Equation 14, first derivatives with Equation 22-24, and working variables with Equation 26-28. With working variables, we augment **C** to form

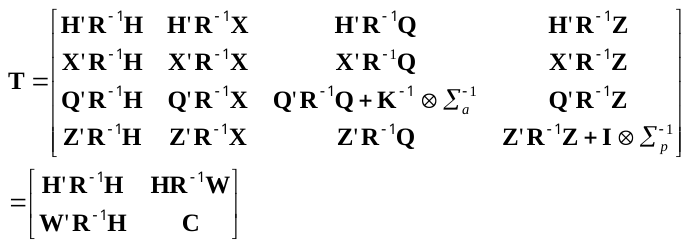
 (34)

Where
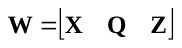
.

Following Gilmour, et al. (1995), matrix **C** is absorbed to construct

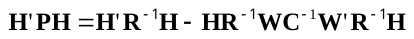
 (35)

Here, the *i*th column of
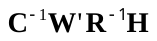
 can be obtained by solving the mixed model equation (Equation 12) with **y** replaced by
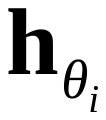
.

Overall, the detailed workflow is described below

Step 1: Initial values assigned to parameter vector
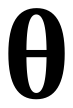
;

Step 2: Build the mixed model equation with Equation 12;

Step 3: Solve the mixed model equation to obtain
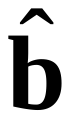
,
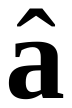
,
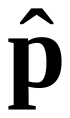
 and
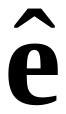
;

Step 4: Calculate first derivatives with Equation 22-24;

Step 5: Calculate the AI matrix with Formula Equation 26-28, 35 and 21;

Step 6: Try AI algorithm with Equation 17, and If the variances go outside the parameter space, implement combined EM and AI algorithm with Equation 33;

Step 7: Repeat the step 2-6 until parameters converge.

2.1.7 **Unbalanced Longitudinal GWAS**

With variances estimated in the null model, we applied two methods to GWAS of unbalanced longitudinal data.

a. **Eigen decomposition of phenotypic covariance matrix**

This method has been described in our previous published paper (Ning*, et al.*, 2018), we directly decompose phenotypic covariance matrix here instead of combined additive and permanent covariance matrix. We detail the theory below.

An additional fixed regression term was incorporated into random regression model to explain the time-dependent SNP effect,

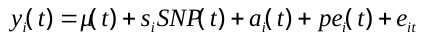
 (36)

Here,
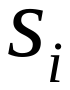
 is a genotype indicator for individual *i* which is assigned 0, 1 and 2 for genotype *aa*, *Aa* and *AA*, respectively; *SNP*(*t*) represents the time-varied additive effect for each marker and can be expressed as linear regression for a set of basis functions as mentioned before,

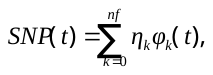
 (37)

where,
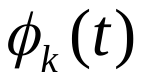
is the value of the *k*th basis function at time *t*;
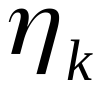
 is the *k*th fixed regression coefficient for additive SNP effect; *nf* is the order of basis functions for the time-varied SNP effect.

The matrix form of (36) can be represented as,

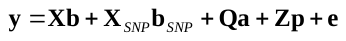
 (38)

Where, is a vector of fixed regression coefficients for SNP effect and is corresponding design matrix.

Similar to the study of Kang*, et al.* (2010) and Zhang*, et al.* (2010), we initially estimated the variance components without including the SNP effect in the model (Equation 8), and then these estimates were applied in the model that examines whole-genome association effects of SNPs. With pre-estimated variance components, phenotypic (co)variance matrix is

(39)

The Eigen decomposition of **V** gives

(40)

where **D** is a diagonal matrix containing the eigenvalues and **U** is the matrix of eigenvectors in the order of the corresponding eigenvalues. We rotate Equation 38 with **U**’, and the mixed model can be rewritten as

(41)

If we define , and , then Equation 41 can be abbreviated as

(42)

The phenotypic (co)variance matrix of rotated model is

(43)

As **V*** is a diagonal matrix, Formula 44 can be solved by weighted least squares

(44)

and

(45)

With the estimated values for fixed regression coefficients of additive SNP effects and corresponding covariance matrix, the Wald Chi-squared test for time-dependent SNP effect is

(46)

b. **Linear transformation of time-varied additive genetic effect**

The method is enlightened by the equivalence between genomic best linear unbiased prediction (G-BLUP) and SNP-BLUP which is derived for univariate GWAS (Gualdron Duarte*, et al.*, 2014; Ning*, et al.*, 2018). Here, we expand the equivalence aforementioned to the field of longitudinal GWAS. Detailed derivation are shown below.

The genome-wide additive time-varied SNP effect model is shown below,

(47)

Here, d is the number of SNPs, and is a centralized SNP indicator for the *j*th SNP of individual *i* which is formulated with Formula 11. represents the time-varied additive effect for the *j*th marker and can be expressed as linear regression for a set of basis functions

(48)

With other symbols same to Equation 1-2, the Equation 47 can be detailed as

(49)

We further adjust it to

(50)

The matrix form for the *i*th individual can be represented as

(51)

Where, is a vector of random regression coefficients for *j*th SNP.

We further adjust (51) with Equation 11

(52)

Here, .

The matrix form for the all individuals can be represented as

(53)

Then, we abbreviate the matrix as

(54)

The (co)variance matrixes for random effects are

(55)

Let , where **a** is a vector of cumulative random regression coefficients for all SNP, *i.e.*, a vector of random regression coefficients for additive genetic effects of individual *i*, then

(56)

The (co)variance matrix for **a** is

(57)

Apply Equation (10) to (57), then

(58)

Thus, Equation (56) is equivalent to Equation (8).

Following Henderson (1949), the best linear unbiased predictor (BLUP) for **u** is

(59)

And the BLUP for **a** is

(60)

As invertible, we can obtain

(61)

Apply (61) to (59),

(62)

The (co)variances for is

(63)

Following Henderson (1975),

(64)

Where is the part of Equation (13).

Therefore, the estimated values of random regression coefficients for *j*th SNP and corresponding (co)variances are

(65)

(66)

The Wald Chi-squared test for time-dependent SNP effect is

(67)

2.2 **Balanced Longitudinal Data**

**2.2.1	Random Regression Model**

For balanced longitudinal data, both  and in Equation (7) are the same for all individuals. If we assume , then . If we assume , Equation (8) can be rewritten as

(68)

**2.2.2** **Variance Components Estimation for Balanced Longitudinal Data**

Inspired by the research of Lee and van der Werf (2016), we applied Eigen decomposition of the genomic relationship matrix to expedite the variance components estimation for balanced longitudinal data. We showed our derivation below.

Eigen decomposition of the genomic relationship matrix **K** gives **K = UDU**’. We rotate (68) with, and the mixed model can be rewritten as

(69)

The phenotypic (co)variance matrix is

(70)

In our study, we estimate variance components using direct REML method for balanced longitudinal data. The restricted log-likelihood function can be formulated as

(71)

Where

The first derivative for each variance component is

(72)

(73)

(74)

The element of AI matrix for variance covariance component *ij* and *i’j’* is

(75)

(76)

(77)

(78)

(79)

(80)

The inversion of matrix **EM** corresponding to and , and , as well as and can be obtained from Equation 30-32.

Similar to variance components estimation for unbalanced longitudinal data, combined EM and AI algorithm is used to improve the calculation efficiency.

**2.2.3 Efficient Computation**

We describe in this section the efficient calculations of the restricted log-likelihood function, the first-order partial derivatives with respect to , and the second-order partial derivatives with respect to and . The first-order and second-order partial derivatives with respect to other parameters can be calculated in a similar fashion.

From (70), we can see that **V*** is a matrix with *n* diagonal blocks of size *t*, where *t* is the number of records for each individual. The computational complexity for obtaining the inverse and determinant of **V*** is only O(*nt*^3^).

For , we have

(81)

The product of **P^*^** and **y^*^** can be efficiently obtained as

(82)

Where

, (83)

(84)

and (85)

For term,

(86)

If we replace **y*** with in (82), the term can be efficiently obtained similarly. Given , and , the vector-matrix-vector product terms for the restricted log-likelihood function, the first-order partial derivatives and the second-order partial derivatives can be calculated efficiently by vector-vector multiplication.

For the trace term in the first derivative, *i.e.*, , we have

Therefore, it involves only the diagonal part matrix multiplication.

**2.2.4 Balanced Longitudinal GWAS**

**a. Eigen decomposition of genomic relationship matrix**

Similar to unbalanced longitudinal GWAS, an additional fixed regression term was incorporated into random regression model to explain the time-dependent SNP effect. We will estimate variance components efficiently for each SNP benefited from the Eigen decomposition of genomic relationship matrix. As most of SNPs contribute little to phenotypes, we considered the estimated variances in the null model as the intimal values for each SNP. This will significantly reduce the interaction times.

b. **Linear transformation of time-varied additive genetic effect**

Similar to Equation (59) for unbalanced longitudinal GWAS, we have

(87)

The (co)variance matrix of estimated SNP effects is

(88)

We can prove that

(89)

Therefore, Equation (88) can be further simplified

(90)

For the *j*th SNP, the estimated effect and corresponding variance are

(91)

(92)

The Wald Chi-squared test for time-dependent SNP effect is

(93)

**References**

Gilmour, A.R., Thompson, R. and Cullis, B.R. (1995) Average information REML: an efficient algorithm for variance parameter estimation in linear mixed models, *Biometrics*, 1440-1450.

Gualdron Duarte, J.L.*, et al.* (2014) Rapid screening for phenotype-genotype associations by linear transformations of genomic evaluations, *BMC bioinformatics*, **15**, 246.

Henderson, C. (1949) Estimation of changes in herd environment, *J. Dairy Sci*, **32**, 706715.

Henderson, C.R. (1975) Best linear unbiased estimation and prediction under a selection model, *Biometrics*, 423-447.

Jensen, J.*, et al.* (1997) Residual maximum likelihood estimation of (co) variance components in multivariate mixed linear models using average information.

Johnson, D. and Thompson, R. (1995) Restricted maximum likelihood estimation of variance components for univariate animal models using sparse matrix techniques and average information, *Journal of dairy science*, **78**, 449-456.

Kang, H.M.*, et al.* (2010) Variance component model to account for sample structure in genome-wide association studies, *Nature genetics*, **42**, 348-354.

Lee, S.H. and van der Werf, J.H. (2006) An efficient variance component approach implementing an average information REML suitable for combined LD and linkage mapping with a general complex pedigree, *Genetics, selection, evolution : GSE*, **38**, 25-43.

Lee, S.H. and van der Werf, J.H. (2016) MTG2: an efficient algorithm for multivariate linear mixed model analysis based on genomic information, *Bioinformatics*, **32**, 1420-1422.

Mrode, R.A. (2014) *Linear models for the prediction of animal breeding values*. Cabi.

Ning, C.*, et al.* (2018) A rapid epistatic mixed-model association analysis by linear retransformations of genomic estimated values, *Bioinformatics*, **34**, 1817-1825.

Ning, C.*, et al.* (2018) Eigen decomposition expedites longitudinal genome-wide association studies for milk production traits in Chinese Holstein, *Genetics, selection, evolution : GSE*, **50**, 12.

VanRaden, P.M. (2008) Efficient methods to compute genomic predictions, *Journal of dairy science*, **91**, 4414-4423.

Verbyla, A.P. (1990) A conditional derivation of residual maximum likelihood, *Australian & New Zealand Journal of Statistics*, **32**, 227-230.

Zhang, Z.*, et al.* (2010) Mixed linear model approach adapted for genome-wide association studies, *Nature genetics*, **42**, 355-360.
